# Benchmarking Twist Genotyping-by-Sequencing Against Whole-Genome Sequencing in Nuclear Families

**DOI:** 10.64898/2026.07.31.742127

**Authors:** Jacob Klugerman, Ivan Iossifov, Kenny Ye

## Abstract

Genome-wide genotyping is widely used in human genetics research, and targeted sequencing-based approaches such as the Twist Bioscience genome-wide SNP capture platform (GxS) have emerged as alternatives to conventional SNP arrays. Here, we evaluated GxS genotype calls from 555 individuals in 184 nuclear families against matched whole-genome sequencing (WGS) calls and compared platform performance with that of the Illumina Infinium Global Screening Array-24 (GSA), which was evaluated in 987 individuals from 279 nuclear families. Genotype data were harmonized across platforms, and analyses were restricted to overlapping SNP loci. Across all callable positions, mean per-SNP call rates were 98.31% for GxS and 98.77% for GSA. Overall SNP concordance with WGS was 99.78% for GxS and 99.66% for GSA. Mean per-individual concordance rates for GxS and GSA were matching their counterpart overall concordance rates when rounded to the nearest hundredth. Per-trio Mendelian violation rates of GxS are about 32 times those of WGS, while those of GSA are about 3.6 times those of WGS on average. These results indicate that GxS performs slightly better than GSA by key concordance metrics, while slightly faltering behind GSA in inheritance analyses.

## Background

Genome-wide genotyping is widely used in human genetics research, including genome-wide association studies (GWAS) and polygenic risk prediction. SNP array platforms such as the Illumina Infinium Global Screening Array-24 have been widely adopted for their low cost, high reproducibility, and established analytical workflows. Combined with genotype imputation, SNP arrays can capture a large proportion of common human genetic variation [1,2].

More recently, targeted sequencing-based genotyping approaches have emerged as alternatives to conventional SNP arrays. The Twist Bioscience genome-wide SNP capture (GxS) platform uses hybridization-based enrichment to interrogate genome-wide SNP loci and may offer advantages in assay flexibility and compatibility with sequencing-based workflows. A recent study demonstrated the utility of the Twist GxS platform for genotyping challenging and degraded DNA samples [4].

However, we were unable to find a third-party evaluation of the Twist Bioscience SNP capture platform or any similar platforms in the existing literature. In this study, we evaluate genotypes called by the Twist Bioscience genome-wide SNP capture platform (GxS) on 555 individuals comprised of 184 nuclear families, with genotype calls by whole-genome sequencing (WGS) [3] on the same individuals serving as a benchmark. Moreover, we compare performance metrics of GxS, such as genotype concordance with WGS and Mendelian violation frequency, to those of the Illumina GSA-24 platform (GSA), which are evaluated similarly on 987 individuals of 279 nuclear families, with no overlap with the 555 individuals genotyped by GxS.

## Methods

### Study Design and Data Sources

This study used genetic data from the SPARK cohort [5], including whole-genome sequencing (WGS) data from SPARK iWGS release v1.1 [6] and whole-exome sequencing (WES) data from SPARK iWES release 3 [7]. Within the iWES release, samples from Batches 1–4 were additionally genotyped using Illumina Global Screening Array version 1 (GSAv1) or version 2 (GSAv2), whereas samples from Batches 5–9 were genotyped using the Twist Bioscience Genomic Screening (GxS) platform. Among subjects genotyped using the GxS platform, 184 nuclear families with a total of 555 individuals were also included in the iWGS release. Among subjects genotyped using the GSAv1 or GSAv2 platforms, 279 nuclear families with a total of 987 individuals were also included in the iWGS release. The performance of GxS was evaluated using WGS calls as a benchmark on these nuclear families and compared to the performance of GSA, which was evaluated in the same way.

### Genotype harmonization

Genotype data generated from the Twist Bioscience genome-wide SNP capture platform (GxS) and the Illumina Infinium Global Screening Array-24 (GSA) were compared against whole-genome sequencing (WGS) genotype calls. Note that in both platforms, only SNP markers excluding indels were targeted. For GSA, only those SNP markers present in both GSAv1 and GSAv2 platforms were retained in our analysis.

Using PLINK v2.0 [8], GxS and GSA genotype data in binary PLINK format included in the iWES release 3 were converted into PED/MAP text formats. WGS genotype calls of iWGS provided in the variant call format (.vcf) were also converted into PED/MAP formats to facilitate comparison across platforms. Multi-allelic positions in the WGS genotype calls were first split to bi-allelic format using bcftools (version 1.10.2) [9] the normalized positions and alleles of the split bi-allelic variants are extracted from the third column (ID) of the vcf file. Genotype harmonization was performed by matching SNP loci of GxS (or GSA) to WGS called variants on chromosome positions and two alleles. SNPs on the GxS (or GSA) platform that do not have a matched variant in WGS genotyping calls were not retained for downstream analyses.

### Concordance and Call Rates

For each SNP in GxS (or GSA) that had a WGS variant matched by exact position and matched alternative alleles, we compared the genotype calls on each individual directly where both platforms made eligible SNP calls. Insertions, deletions, and other non-single-base variants called by WGS were treated as missing, as well as no-calls made by either WGS or the platform being evaluated. Matching genotypes were classified as concordant, and all others as discordant. Concordance and call rates were aggregated across all SNP positions and individuals. We further classified concordance rates by six SNP variant types (A/C, A/G, A/T, C/G, C/T, G/T). To assess systematic genotype-specific error patterns between the tested platforms and WGS, genotype confusion matrices were generated for each of six variant types.

### Mendelian violations

Mendelian inheritance consistency was evaluated using parent–child trios available within each dataset. Mendelian violations were checked only on SNP loci for which genotype calls were available for the child and both parents in both the tested platform and WGS. Mendelian violation rates were aggregated for each trio and normalized by the number of eligible loci.

### Filtering by concordance and call rate

The performances of GxS and GSA were re-evaluated on a subset of SNPs that possessed at least 99% call rates across all platforms and at least 99% concordance rates with WGS.

## Results

### Call rates

All metrics reported in this section are summarized in Table 1. Across all positions in both datasets, the mean per-SNP call rate, aggregated over all individuals, was 98.31% for GxS, with a median of 100% and a 10^th^ percentile of 98.02%. In comparison, the mean, median, and 10^th^ percentile rates for GSA were 98.77%, 99.80%, and 97.26%, respectively. In GxS, 55.70% of positions had a call rate of 100%, and 85.24% had a call rate of at least 99%. In GSA, 28.44% of positions had a call rate of 100%, and 78.55% had a call rate of at least 99%.

**Table 1.** Call rate, concordance, and Mendelian violation metrics for GxS and GSA relative to WGS, before and after filtering. Call rates are computed over all individuals genotyped on each platform; concordance is computed over genotype pairs in which both WGS and the respective platform produced a call; Mendelian violation rates are means per trio.

|  | GxS | GSA |
| --- | --- | --- |
| **Cohort** |  |  |
| Individuals genotyped | 555 | 987 |
| Mother-father-child trios | 187 | 429 |
| Nuclear families | 184 | 279 |
| **Call rate** |  |  |
| All assay markers included. |  |  |
| Variants | 1,380,542 | 589,147<sup>a</sup> |
| By position, mean | 98.31% | 98.77% |
| By position, $\geq 99\%$ | 85.24% | 78.55% |
| By individual, mean | 98.26% | 98.67% |
| By individual, $\geq 99\%$ | 21.80% | 48.94% |
| **Concordance with WGS** |  |  |
| Only including genotype pairs called in both WGS and the platform. |  |  |
| Matched variants | 1,379,968 | 571,358 |
| By position, mean | 99.77% | 99.63% |
| By position, $\geq 99\%$ | 95.44% | 96.82% |
| By individual, mean | 99.78% | 99.66% |
| By individual, $\geq 99\%$ | 100.00% | 98.78% |
| Overall | 99.78% | 99.66% |
| Overall, after filtering | 99.92% | 99.99% |
| **Mendelian violation rate** |  |  |
| Mean per-trio rate computed over matched markers with calls in each member of the trio. |  |  |
| Genotyping assay | 0.1093% | 0.0354% |
| Genotyping assay, after filtering | 0.0477% | 0.0055% |
| WGS (Matched assay markers) | 0.0034% | 0.0098% |

a. Markers present on both GSAv1 and GSAv2.

The mean and median per-individual call rates were 98.26% and 98.53% for GxS, and 98.67% and 98.99% for GSA. Among 555 individuals genotyped on GxS, 121 (21.80%) exhibited call rates exceeding 99% across all positions; out of 987 individuals genotyped on GSA, 483 (48.94%) had call rates greater than 99%. On the other hand, only 1 out of 555 (0.18%) GxS individuals had a call rate below 95%, while 18 out of 987 (1.82%) GSA individuals did.

### Concordance with WGS genotyping Calls

Out of 1,380,542 GxS SNP markers, 1,379,968 were matched to WGS variants by their positions and alleles. Of 589,147 GSA SNP markers present in both GSAv1 and GSAv2 platforms, 571,358 were matched to WGS variants. The overall concordance rates across all individuals and all positions for GxS and GSA were 99.78% and 99.66%, respectively. Concordance rates varied by variant type (Fig. 1a), with variant A/T having the lowest concordance rate in both GxS and GSA at 99.72% and 99.28% respectively.

**Figure 1.**
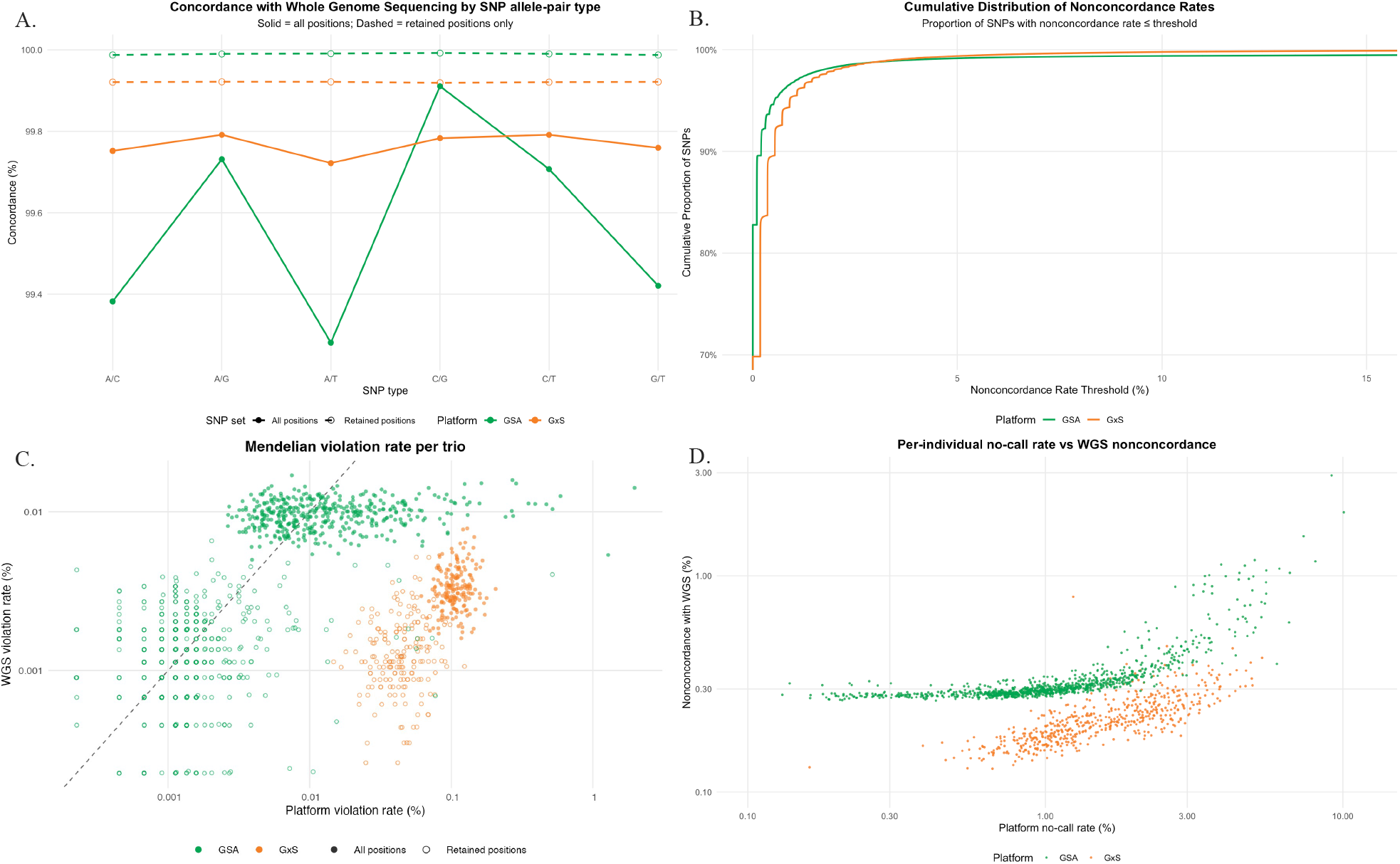
Comparison of genotype performance metrics between Twist Genotyping-By-Sequencing (GxS) and the Global Screening Array (GSA) using whole-genome sequencing (WGS) as the reference standard. **(A)** Genotype call concordance rates for GxS and GSA relative to WGS, stratified by SNP position type based on unordered reference/alternate (REF/ALT) allele pairs. **(B)** Cumulative distribution of per-SNP non-concordance rates for GxS and GSA compared with WGS. **(C)** Per-family Mendelian violation rates for GxS and GSA relative to WGS across all analyzed variant positions and across the subset of positions with genotype call rates ≥99%. **(D)** Per-individual no-call rates and per-individual non-concordance rates with WGS for GxS and GSA.

The cumulative density plots of non-concordance rates over all positions, shown in Fig. 1b, detail that GSA has more high-concordance positions than GxS, yet falls slightly below GxS at higher non-concordance thresholds due to a nontrivial number of positions at lower concordance rates. A total of 95.44% of GxS positions had a non-concordance rate below 1% compared to 96.82% of GSA positions, but a total of 99.38% of GxS positions had a non-concordance rate below 5% compared to GSA’s 99.16%.

The mean per-individual concordance rates with WGS were 99.78% in GxS and 99.66% in GSA, with medians of 99.79% and 99.70%, respectively. All individuals in GxS and 975 out of 987 individuals in GSA had genotype concordance rates greater than 99%. As shown in Figure 1d, per-individual non-concordance rates were positively associated with per-individual no-call rates among both GxS and GSA. The relationship is non-linear but tight, with Spearman correlations of 0.81 for GxS and 0.85 for GSA. Moreover, it is also clear from the figure that GxS per-individual call-rates and concordance rates show less variation than those of GSA.

### Mendelian violations

There are 187 trios from 184 nuclear families for GxS, and 429 trios from 279 nuclear families for GSA. Among the 187 mother-father-child trios present in GxS data, the average Mendelian violation rate across all valid positions was 0.1093%. In contrast, the Mendelian violation rate was 0.0034% in matched WGS calls, more than 32 times lower. The average Mendelian violation rate in GSA was 0.0354% across 429 family trios and 0.0098% in matched WGS calls. These observations confirm that WGS genotyping calls are superior to those from either GxS or GSA.

As shown in Figure 1c, GxS mendelian violation rates for each parent-child trio were consistently above the corresponding rates of WGS, whereas the rates of GSA showed greater variability across trios, and in many cases were on par or even lower than those of WGS.

### Filtering

In practice, bad SNP markers, often flagged by low call rates are removed from further analysis. After filtering out SNP markers with concordance rates are less than 99% and calls rates are less than 99% of individuals by either the platform being evaluated or WGS, 1,146,589 positions were retained for GxS and 444,495 for GSA. The concordance rates with WGS for the retained SNPs increased to 99.92% for GxS and 99.99% for GSA. The concordance rates also become very consistent across the six variant types (Fig. 1a).

Per-family Mendelian violation rates for GxS, GSA, and WGS were recalculated after the filtering. Among the 187 trios in GxS, the Mendelian violation rate for retained positions decreased by a factor of 2.3 to 0.0477%, and in GSA, the per-family Mendelian violation rate decreased by a factor of 6.4 to 0.0055%. However, the per-family rates of GxS remained consistently higher than the corresponding WGS rates (Fig. 1c).

## Discussion

This study is motivated by the lack of published third-party studies on Twist GxS platforms, while we took advantage of overlapped samples in SPARK iWESv3 release and iWGSv1 release.

To evaluate the quality of GxS genotyping calls, we matched variants on GxS (and GSA) within the WGS variant calls by position and allele. A very small and negligible proportion (0.04%) of GxS markers were not matched to WGS variant calls, but we found the number of unmatched GSA SNP markers were much greater at almost 3%, for which we will address later.

For the matched positions, since GxS and GSA only call SNPs by design, we removed all WGS indel calls from the comparison. Removed indel calls represent less than 0.3% of total calls at these positions, and their number is less than 20% of the number of no calls by WGS.

Our analysis showed that GxS has a slightly better concordance rate than GSA on average. However, these rates were aggregated over largely different SNP markers. Therefore, we compared average concordance rates for the 75,634 variants that were present in both platforms, stratifying by six variant types. As shown in Supplementary Figure 1, GxS performs slightly better than GSA at these overlapped positions. The slight advantage of GxS mainly come from multi-allelic positions (as seen in WGS), on which the concordance between GSA and WGS reduces to 95.3% while the concordance between GxS and WGS stays at 99.6%. Taking out the multi-allelic positions, the concordance between GxS and WGS is actually slightly lower than that between GSA and WGS.

We further investigated discordances between GxS and WGS and the discordances between GSA and WGS. Table 2 shows confusion matrices for each variant type. As can be seen, GxS is much more likely to incorrectly call heterozygous genotypes homozygous than GSA, but also much less likely to call homozygous genotypes heterozygous or homozygous of the opposite allele. This difference can be explained by the different technology used by these two platforms. GxS is sequence-based, so if one of two alleles at a heterozygous site is not observed, a homozygous call is made. The chance of this happening is 1/2^n-1^, where *n* is the number of reads observed at this site. At n=10, this is approximately .2%. This vulnerability also explains the high Mendelian violation rate observed in GxS calls, as it occurs when a heterozygous genotype of a parent is called homozygous.

**Table 2.**
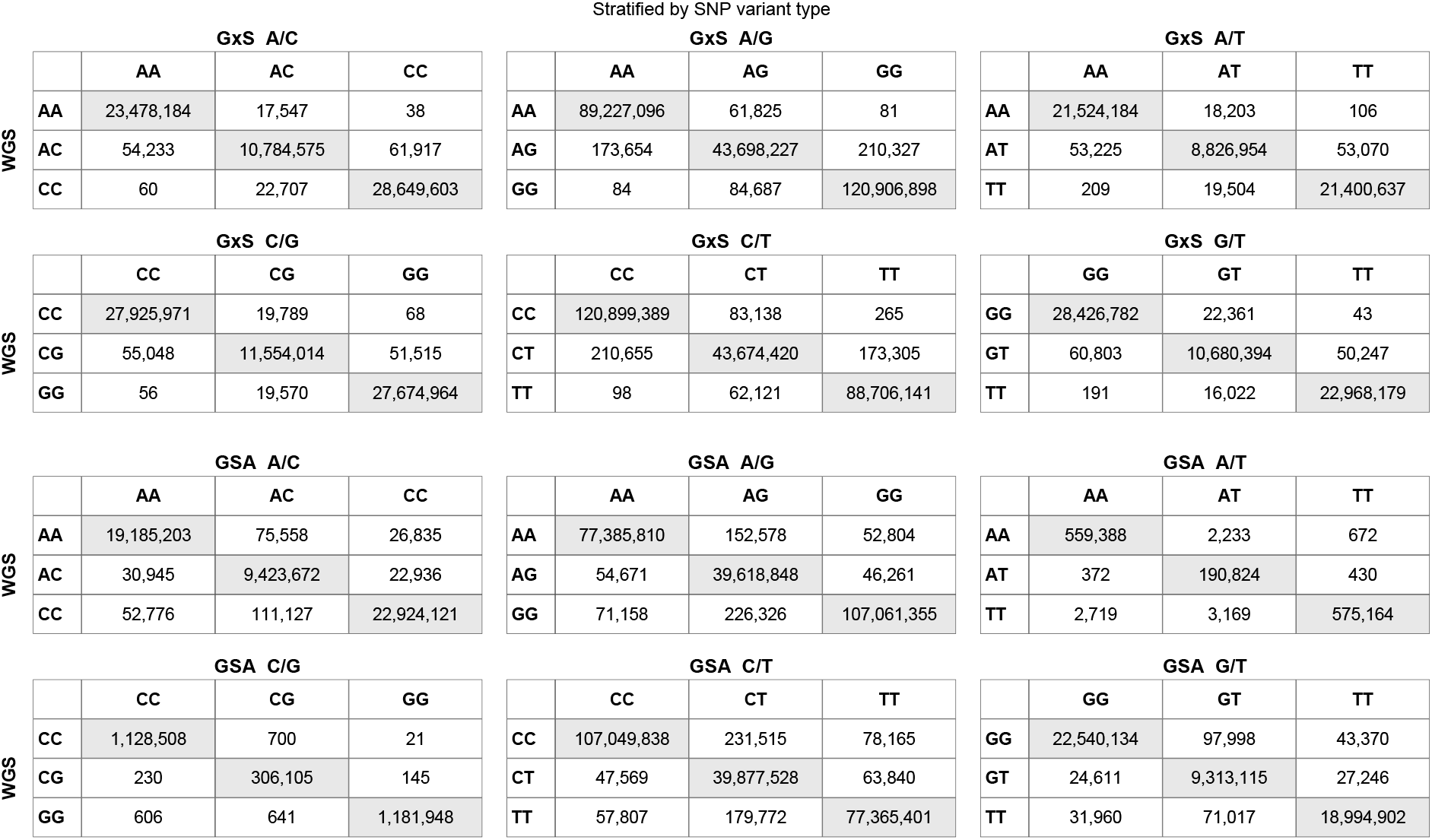
Confusion matrices of genotype calls for GxS and GSA relative to WGS, stratified by SNP variant type. Matching variant types are shaded grey.

As mentioned earlier, as much as 3% of GSA SNPs were not matched to WGS variants. A large majority of them can be explained by being rare in population. According to Allele Frequency Aggregator (ALFA), (*www.ncbi.nlm.nih.gov/snp/docs/gsr/alfa/*), 95% of these unmatched SNPs having MAF < 0.001 and 82% < 0.0001, therefore they may not be present in the iWGS cohort. However, more than 300 unmatched SNPs, according to ALFA, have MAF > 0.1, yet none of them were reported by iWGSv1.1, which includes 6756 founders. We further noticed that for some of those SNPs, there is a large discrepancy in reported allele frequencies between ALFA and other databases, notably, GnomAD. For example, for rs77580625 (chr1:76759287,C>T), ALFA reported the frequency of T allele being 0.51 while GnomAD reported 0.00002. The latter is consistent with SPARK iWGS release. While this is out of scope of this study, it might worth further investigation on the source of such discrepancies. Either some of the data ALFA used were outdated, or the genotyping call pipeline used by iWGS has some deficiencies. For this purpose, we included a list of unmatched GSA SNP markers in the supplementary, as well as their allele frequencies reported by ALFA.

## Conclusion

Overall, we found that the Twist GxS genotyping platform is highly accurate. Its concordance with WGS is slightly higher than the more traditional Illumina array platform. However, as it subject to a higher error rate of calling heterozygous genotype homozygous, it tends to generate higher mendelian violation error. In addition, compared to GSA, GxS has lower assay-to-assay variation, as per-individual call rates and concordance rates are more consistent. Nonetheless, we also observed a larger share of GxS than GSA positions with concordance rate and call rate below 99%, and we recommend that those positions be filtered out for genetic analysis.

## Ethics approval and consent to participate

All data were obtained from the SPARK cohort under an institutional data access agreement. The original SPARK study received IRB approval from SFARI. No additional ethics approval was required for this secondary analysis.

## Consent for publication

Not applicable.

## Availability of data and materials

The datasets analyzed in this study are available from the Simons Foundation Autism Research Initiative (SFARI) upon approved application at https://base.sfari.org. The individual ID and family ID of the subjects included in the study are provided in the Supplementary materials in the PLINK .fam format. Computer programming code is available from the first author upon reasonable request.

## Authors’ contributions

JK performed data analysis and drafted the manuscript. II contributed to the conception and design of the study. KY conceived and designed the study and supervised the data analyses. All authors read and approved the final manuscript.

## Acknowledgements

The authors acknowledge using OpenAI’s ChatGPT (GPT-5.5) to generate an initial draft of portions of the introduction. The authors reviewed, edited, and revised all generated text and take full responsibility for the accuracy and integrity of the final manuscript. We are grateful to all the families at the participating SPARK sites as well as the SPARK Consortium for contributing the genetic data that made this research possible. We appreciate the opportunity to access phenotypic and genomic sequencing datasets (including SPARK iWGS release v1.1 and iWES release 3) on SFARI Base. We thank the Simons Foundation Autism Research Initiative (SFARI) for establishing and funding the SPARK cohort [5].

---

**Supplementary Figure 1.**
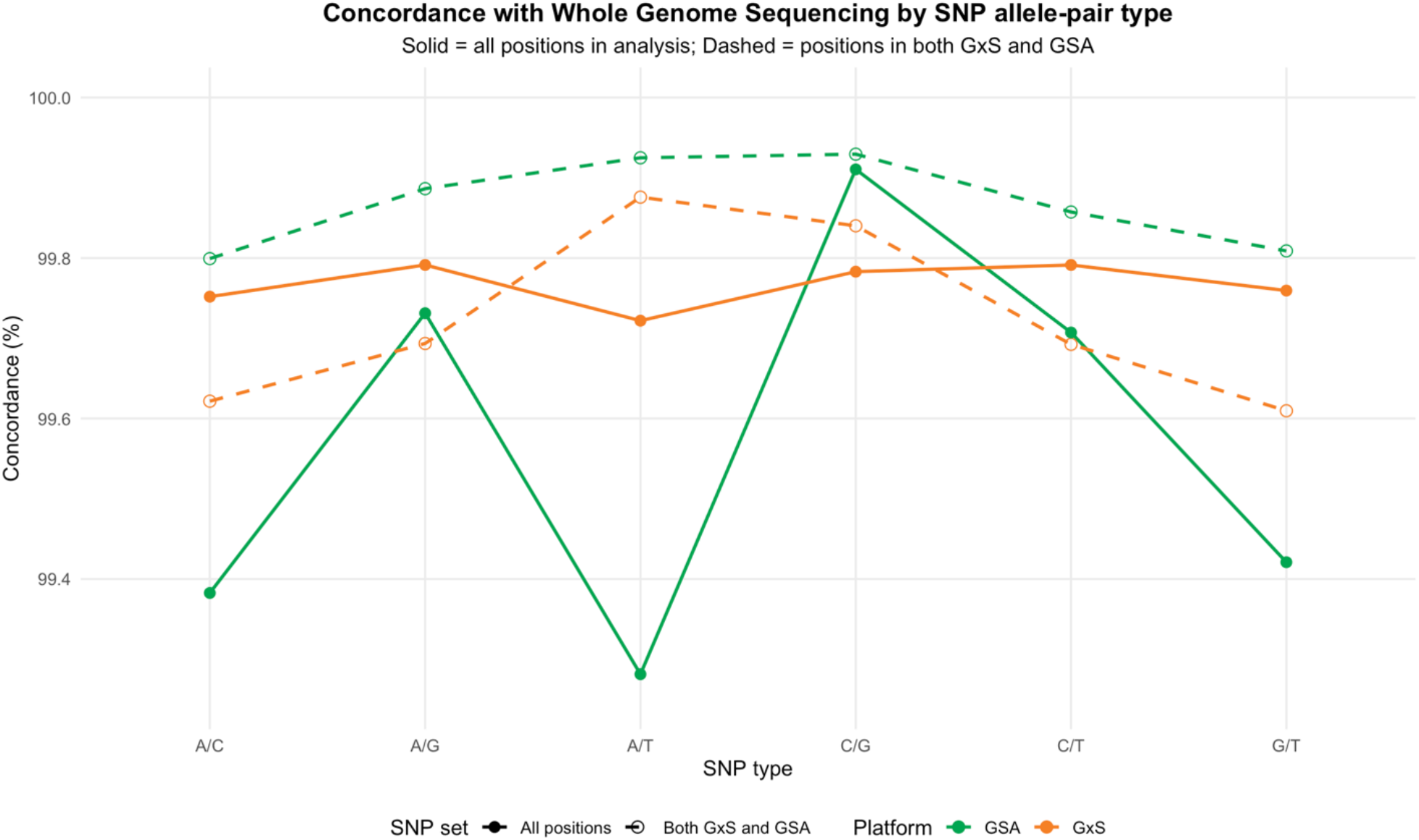
Per-position concordance rates with Whole Genome Sequencing by SNP allele-pair type including overlapping SNP positions between Twist Genotyping-By-Sequencing (GxS) and Illumina Global Screening Array (GSA). The dashed line represents concordance rates for positions that were present in both GxS and GSA comparisons with WGS.

